# Mutation-induced heterogeneity of the β7-β8 loop of the *Staphylococcus aureus* class A sortase leading to enhanced catalytic efficiency characterized by NMR and enzyme kinetics

**DOI:** 10.64898/2026.08.21.746310

**Authors:** Erich G. Walkenhauer, Noah Cox-Tigre, Manish Chaubey, Raphael Marcenac, Adam Wachsman, Hanna M. Kodama, Katy M. Lindblom, Claire E. Bloom, John M. Antos, George P. Lisi, Serge L. Smirnov, Jeanine F. Amacher

**Affiliations:** Department of Chemistry, Western Washington University, Bellingham, WA, USA; Department of Molecular Biology, Cell Biology & Biochemistry, Brown University, Providence, RI, USA; Department of Biology, Whitman College, Walla Walla, WA, USA; Giuliani RNA Center, Brown University, Providence, RI, USA

**Keywords:** sortases, enzymes, conformational states, structural biology, solution NMR, specificity

## Abstract

Bacterial sortase enzymes are cysteine transpeptidases at the surface of Gram-positive bacteria that ligate substrates to the cell wall. In addition, these enzymes are powerful tools in protein engineering applications *via* sortase-mediated ligation (SML) due to their covalent attachment of two substrates, with one containing a pentapeptide recognition motif with sequence LPXTG, where X=any amino acid, and the second, an N-terminal glycine. The class A sortase from *Staphylococcus aureus* (saSrtA) was the first to be identified, and over 25 years later, the most widely used SML variants continue to be derivatives of a directed-evolution-identified pentamutant of saSrtA, or saSrtA5M. We previously characterized P94, a position mutated in saSrtA5M that interacts directly with a structurally conserved loop (the β7-β8 loop) near the active site of wild-type saSrtA only in the inactive conformation. This work revealed that the single P94X mutation dramatically affects relative saSrtA activity, as well as specificity for the P2 (or X) position in the LPXTG recognition motif. This is largely driven by *K*_m_ effects. Here, we further interrogated P94 by probing structural changes in the active, apo state of saSrtA in the presence of the P94D mutation, as well as via mutations in Y187, the β7-β8 loop residue hypothesized to interact directly with P94. The saSrtA enzyme is allosterically activated by calcium; therefore, we were interested if P94D would induce structural changes in the calcium-bound apo enzyme. We used ^1H^-^15^N NMR experiments to compare spectra between enzymatically inactive variants of saSrtA with and without the P94D mutation. We also used NMR to calculate relative binding affinities for a pentapeptide substrate to these variants, as well as enzymatically inactive saSrtA5M. Our NMR data, in combination with enzymatic assays using active variants confirmed differences in the active, apo states of these enzymes. Overall, this work provides additional atomic detail regarding the importance of the P94 residue in saSrtA substrate recognition.

## Introduction

Bacterial sortases are cysteine transpeptidases at the cell surface of Gram-positive bacteria, e.g., of *Staphylococcus* and *Streptococcus* genera. First identified in 1999 in *Staphylococcus aureus*, class A sortases (SrtAs) are generally considered to be ‘housekeeping’ enzymes, and are responsible for recognizing and tethering substrate proteins to the growing peptidoglycan cell wall.^1–4^ Substrates include bacterial toxins, environmental sensors, and other critical proteins.^3,4^ There are multiple recognized classes of sortases (A-F) which are responsible for differing activities, e.g., heme transport (class B sortases) or pilus biogenesis (class C sortases).^4^ Class A sortases remain the most well studied. Due to their ability to ligate two sequences together, these enzymes are also widely utilized in protein engineering applications via sortase-mediated ligation (SML) applications.^5–7^ For SML experiments, *S. aureus* SrtA (saSrtA) derivatives remain the most widely used.^2,6^

The SrtA mechanism is well studied. SrtA enzymes recognize substrate proteins using a Cell Wall Sorting Signal, defined as the sequence L-P-X-T-G where X=any amino acid; notably, relative promiscuity for variations in this sortase recognition motif differs amongst SrtA enzymes.^3,8–11^ All sortases share a conserved catalytic triad, consisting of Cys-His-Arg residues, though the catalytic role of the arginine residue is debated. Recent data from our group and others suggests that a Thr residue immediately preceding the catalytic Cys, along with the backbone amide of the residue immediately following the catalytic His stabilize the generated oxyanion intermediates, with the Arg playing a stabilizing role for the bound ligand.^2,12–14^ Nucleophilic attack by the thiol group of the catalytic Cys on the carbonyl carbon between the Thr (termed P1) and Gly (P1’) residues generates an oxyanion intermediate, followed by the formation of an acyl enzyme adduct and release of the P1’ Gly plus the C-terminus of the initial LPXTG substrate.^1–3,15^ This is a ping-pong reaction, and nucleophilic attack by an incoming N-terminal glycine-containing second substrate on the P1 carbonyl carbon forms the second tetrahedral oxyanion intermediate, which is subsequently resolved with product release.^2,3^ Ultimately, the resulting ligated product includes the N-terminal portion (LPXT-) of the first substrate and entirety of the second substrate (**Figure 1A**).

Despite its potential as a protein engineering tool, there were early challenges with utilization of saSrtA in SML experiments, namely, its strict specificity for the LPXTG substrate sequence, relatively low catalytic efficiency (200 M^-1^ sec^-1^ for an LPETG peptide), off-target hydrolysis product formation, and reaction reversibility.^2,5,6^ There have been notable advances in modulating target specificity, e.g., by using evolved versions of saSrtA or other SrtA enzymes with natural variations in recognition, and reducing reaction reversibility, e.g., by using metal sequestration or electrostatically assisted capture of specific substrates and/or enzyme-catalyzed degradation.^16–22^ Of these, a major advance in the SML field was the development of a pentamutant via directed evolution, with the reported saSrtA5M variant (P94R/D160N/D165A/K190E/K196T) having a *k*_cat_/*K*_m_ = 23,000 M^-1^ sec^-1^ for an LPETG peptide, a 115-fold improvement in catalytic efficiency. This result was largely driven by a 33-fold reduction in *K*_m_ (7.6 mM in wild-type (WT) versus 0.23 mM in saSrtA5M), and further enhanced by a 3.6-fold increase in *k*_cat_ (1.5 s^-1^ in WT versus 5.4 s^-1^ in saSrtA5M).^16^

Despite widespread use of saSrtA5M, and a related heptamutant (saSrtA7M) with two added mutations (E105K/E108A or E105K/E108Q)^23–25^ that remove a dependence on calcium for activation, the stereochemical basis of heightened activity conferred by the saSrtA5M mutations is not known. SrtA enzymes share a general antiparallel 8-stranded β-barrel ‘sortase-fold’ core, with varying numbers of α-helices and loops connecting the β-strands (**Figure 1B**).^2^ Comparison of previously reported experimental structures of saSrtA in the inactive, apo (+Ca^2+^) and ligand-bound (+Ca^2+^) forms (PDBs 1IJA and 2KID)^26,27^ revealed the largest conformational changes are within two structurally conserved loops near the enzyme active site, the β6-β7 and β7-β8 loops (**Figure 1C**). Extensive investigation of these loops in SrtA enzymes by our group and others previously characterized their overall contribution to substrate recognition and overall activity.^10–12,28,29^ In the structures of saSrtA, specifically, the β7-β8 loop transitions from a downward ‘closed’ position occluding the active site in the apo structure (PDB 1IJA) to an upward ‘open’ position in the active (+Ca^2+^), ligand-bound structure (PDB 2KID) (**Figure 1C**). Interestingly, while the 1IJA structure was solved in the presence of calcium, the NMR structure does not show the fully activated enzyme conformation, indicated by an ordered β6-β7 loop and opening of the β7-β8 loop (**Figure 1C**). Additional work suggested that while the addition of calcium facilitates these structural changes, there are likely intermediate states prior to substrate binding.^27,30^ Ultimately, these structures suggested that the closed conformation is facilitated by a hydrophobic interaction between the side chain of Y187 in the β7-β8 loop and the side chain of P94 in α1 of saSrtA (**Figure 1D**). Interestingly, P94 is one of the mutated amino acids in the saSrtA5M pentamutant.

**Figure 1.**
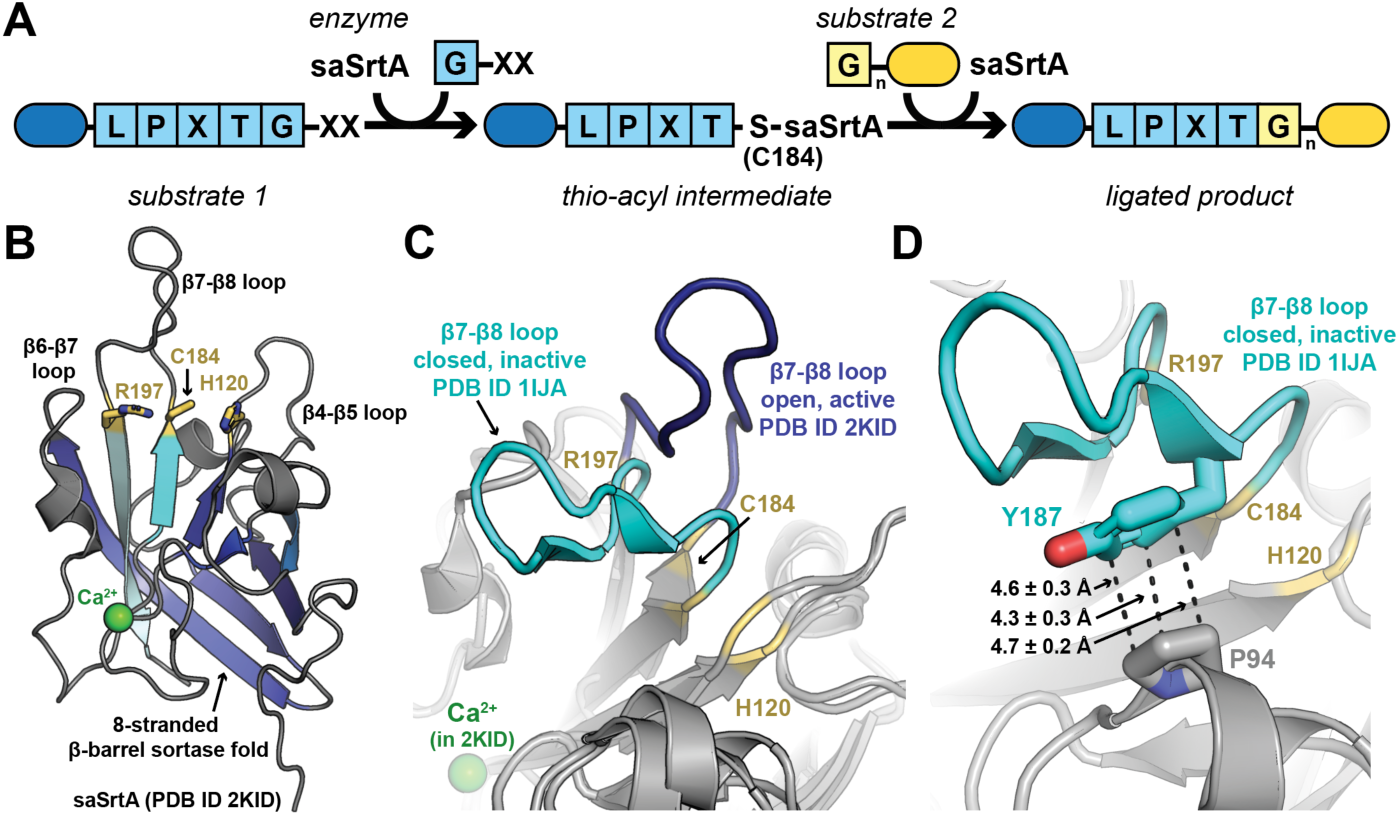
Overall reaction for saSrtA and conformational changes upon activation. (**A**) Reaction schematic for sortase-mediated ligation. (**B**) The conserved 8-stranded anti-parallel β-barrel sortase fold is highlighted on an experimental saSrtA structure (PDB ID 2KID). Here, the structure was determined in the presence of Ca^2+^ and a LPAT* peptidomimetic. The side chains atoms of active site residues are shown as sticks, labeled, and colored by heteroatom (C=yellow, N=blue, S=gold). Structurally conserved loops near the active site (β4-β5, β6-β7, and β7-β8) that are known to affect overall enzyme activity and specificity are labeled. (**C**) The differing closed, inactive and open, active conformational states of the β7-β8 loop in saSrtA are highlighted. Both experimentally determined structures (PDB IDs 1IJA, 2KID) are shown as cartoon, with the loops colored as labeled, and positions of catalytic residues in yellow. The Ca^2+^ ion (2KID) is a green sphere. (**D**) An observed interaction between P94 and Y187 (in the β7-β8 loop) is highlighted in the inactive saSrtA (1IJA) structure. The structure is rendered as in (**C**), but with the side chain atoms of P94 and Y187 shown as sticks and colored by heteroatom (C=gray/cyan, O=red, N=blue). Distances shown are averaged values ± standard deviation as calculated from the 25 NMR states in PDB ID 1IJA.

We recently reported characterization of 19 variants of P94X saSrtA, which included all mutations except for cysteine due to its proximity to the catalytic Cys.^31^ We found that almost all P94X variants were improved over WT, with several, including P94D, also increased as compared to P94R (one of the mutations in saSrtA5M), particularly for substrate sequences without a P2 Glu (LP**E**TG). We were further specifically interested in the P94D variant due to the presence of several negatively charged residues in the β7-β8 loop (D185, D186, and E189), and possibility for repulsive interactions which we reasoned could facilitate the open, active conformation. Enzyme kinetics confirmed that these substitutions predominantly acted by lowering relative *K*_m_ values. In performing these experiments, however, we also wanted to probe differences in the apo (+Ca^2+^) conformations of our saSrtA variants. A recent experimental structure of saSrtA5M in the presence of Ca^2+^ provides insight into the structure of the β7-β8 loop in this evolved variant,^32^ but the dynamics of the closed-to-open transition remain not fully characterized for the WT enzyme.^26,33,34^

To continue to fill this gap, here we used nuclear magnetic resonance (NMR) experiments to characterize conformational differences between WT and P94D saSrtA and to determine the relative binding affinities of inactive C184A variants of these enzymes and saSrtA5M for an LP**A**TG peptide substrate. We further probed the proposed P94-Y187 interaction through mutagenesis of Y187. NMR titration experiments and enzyme kinetics assays demonstrated that the P94D mutation alters substrate affinity (*K*_m_ for active, and *K*_d_ for inactive variants) while maintaining comparable catalytic efficiency for the LP**A**TG substrate. In addition, we identified NMR resonances corresponding to β7-β8 loop residues which differ between WT and P94D saSrtA, suggesting conformational changes upon mutation. Together, these findings provide new structural insight into substrate recognition by saSrtA and related mutants, which we envision will further support its utility for SML technologies.

## Materials and methods

### Protein expression and purification

Recombinant protein expression and purification was performed as previously reported.^10,31,35^ Briefly, WT, P94D, P94D/C184A, C184A, Y187A, Y187D, and Y187R saSrtA variants and saSrtA5M and C184A saSrtA5M sequences were inserted into the pET28a(+) vector (Genscript) with a N-terminal His_6_-tag (HHHHHH) and a Tobacco Etch Virus (TEV) protease cleavage site (ENLYFQ/S) and transformed into *Escherichia coli* BL21 (DE3) cells. All sequences used are included in Supporting Information. Bacterial cells were grown, induced with 0.15 mM isopropyl β-D-1-thiogalactopyranoside (IPTG), and harvested as previously reported.^10,31,35^ For NMR experiments, ^15^N labeled protein expression of C184A saSrtA, P94D/C184A saSrtA, and C184A saSrtA5M was performed by growing transformed BL21 (DE3) cells in LB media until the OD_600_ reached 0.6-0.8. At this point, the growths were spun down at 5,000 x *g* for 10 min at 4°C. The resulting cell pellets were resuspended in 0.5 L of salt wash (8.5 mM NaCl, 22 mM KH_2_PO_4_, 42 mM Na_2_HPO_4_, 4 µM thiamine, 86 µM kanamycin, 1 mM MgSO_4_, 100 µM CaCl_2_) per each 1 L growth. The resuspension was again centrifuged as above, and the resulting cell pellet was resuspended in 1 L of M9 minimal media (salt wash with the addition of 22 mM glucose and 24 mM ^15^NH_4_Cl). The M9 culture was incubated for 1 h at 37°C with shaking at 210 rpm before induction with 0.15 mM IPTG and temperature reduction to 18°C for 18-20 h. Cells were harvested as previously reported.^10,31,35^ For all, the lysis buffer contained 50 mM Tris pH 7.5, 150 mM NaCl, and 0.5 mM ethylenediaminetetraacetic acid (EDTA).

Purification of all proteins was conducted using a 5 mL Ni-NTA column (His-Trap HP, Cytiva) with a wash buffer of 50 mM Tris pH 7.5, 150 mM NaCl, 20 mM imidazole and an elution buffer of the same solution with 300 mM imidazole. The His_6_ purification tag was cleaved using TEV protease for the ^15^N-labeled WT saSrtA, P94D saSrtA, and saSrtA5M proteins, which were dialyzed into a buffer of 50 mM Tris pH 7.5, 150 mM NaCl, 1 mM dithiothreitol (DTT), and 0.5 mM EDTA following elution from the Ni-NTA column. TEV protease was added at 1:14 TEV:protein for 1 h at 34°C with shaking at 100 RPM. The protein was added to a second Ni-NTA column using wash buffer and the flow-through was collected. All proteins were further purified using size exclusion chromatography (HiLoad 16/600 Superdex 75, Cytiva) with gel filtration buffer (50 mM Tris pH 7.5, 150 mM NaCl). SDS-PAGE was used to assess purity at all stages and protein was concentrated using Amicon Ultra-15 Centrifugal Filter Unit (10,000 MWCO) spin concentrators. Protein concentration was determined by measuring A_280_ and using the following theoretical extinction coefficients: 15930 M^-1^ cm^-1^ (WT, P94D, P94D/C184A, and C184A saSrtA variants, and saSrtA5M and C184A saSrtA5M variants) or 14440 M^-1^ cm^-1^ (Y187X saSrtA variants). All proteins were flash frozen using liquid nitrogen and stored at -80°C for assays.

### Peptide synthesis

Solid-state peptide synthesis was performed as previously reported.^10–12,29,31^ The Ac-LP**A**TGG-NH_2_ peptide used in NMR experiments was purchased from Biomatik. The peptide substrates (Abz-LP**X**TGGK(Dnp), with X = A, E, K, S) used for peptide cleavage and enzyme kinetics assays were synthesized using a Biotage Initiator + Alstra peptide synthesizer on Fmoc Rink-Amide MBHA resin on a 0.1 mmol scale as recently described.^31^ Here, Abz = 2-aminobenzoic acid and Dnp = 2,4-dinitrophenyl. All peptides were purified using a Phenomenex Luna 5 μM C18(2) 100 Å column (10 x 250 mm) by reverse-phase HPLC, and their identities were confirmed by mass spectrometry, as previously described.^31^

### Peptide cleavage assays

Activity assays were performed as previously reported.^8,10–12,29,31,35,36^ Briefly, saSrtA variants were tested in reaction buffer (50 mM Tris pH 7.5, 150 mM NaCl, 10 mM CaCl_2_, 5 mM NH_2_OH) in 96-well plates, with a final volume of 100 µL, enzyme concentration of 1 µM, and peptide substrate concentration of 50 µM. Control assays were performed under the same conditions, but lacking the enzyme. All assays were run with [DMSO] < 5% (*v/v*) on a Biotek Synergy H1 plate reader in technical triplicate with λ_excitation_ = 320 nm and λ_emission_ = 420 nm, characteristic of the Abz fluorophore. Fluorescence was recorded every 1 min over a 2 h period using a xenon-flash light source set to high, a read height of 7 mm, and gain set to 75.

### Enzyme kinetics assays

Enzyme kinetics assays were performed as recently reported.^31^ Briefly, the reaction buffer used was 50 mM Tris pH 7.5, 150 mM NaCl, 10 mM CaCl_2_, 1 mM hydroxylamine (H_2_NOH), and a final DMSO concentration of 5% (*v/v*). The saSrtA concentration used was 2.79 μM based on recently published work,^20^ and the Abz-LPATGGK(Dnp) substrate was added at variable concentrations of 10 μM, 25 μM, 50 μM, 100 μM, 250 μM, and 500 μM. The final reaction volume was 200 μL. Assays were run in triplicate and quenched at various initial time points (10, 60, 120, and 180 s) using glacial acetic acid (7:1 final ratio of reaction mixture to acetic acid). The reactions were analyzed using a Dionex Ultimate 3000 HPLC system as previously described.^31^ GraphPad Prism 11 was used to plot the data and calculate *k*_cat_ and *K*_m_ using the Michaelis-Menten equation for each replicate.

### NMR Data Acquisition and Processing

Protein samples were exchanged into NMR buffer (50 mM Tris pH 6.2, 150 mM NaCl, 10 mM CaCl_2_, 3 mM dithiothreitol (DTT), 7% deuterium oxide (D_2_O)) using an Amicon 10k MWCO spin concentrator and then concentrated to 1 mM protein concentration. NMR experiments were performed on an 18.8 T Bruker Avance NEO spectrometer operating at 850.13 MHz ^1^H, 213.76 MHz ^13^C, and 86.18 MHz ^15^N frequencies. All measurements were performed at 298 K. The NMR data were processed with NMRPipe and the resulting NMR spectra were analyzed and visualized with NMRViewJ and Sparky^37–39^. Resonance assignments of WT saSrtA were transferred from Biological Magnetic Resonance Bank (BMRB) entry 4879.^26,40^

NMR titrations for the C184A saSrtA, P94DC184A saSrtA, and C184A saSrtA5M enzymes with Ac-LPATGG-NH_2_ peptides were performed by collecting a series of ^1^H-^15^N HSQC spectra with increasing peptide concentrations until no further spectral perturbations were detected; this included 0-5.5 mM substrate for C184A saSrtA, 0-3.5 mM for P94D/C184A saSrtA, and 0-6.5 mM for C184A saSrtA5M. NMR chemical shift perturbations (CSP) were calculated as Δδ=√((*δ_H_*^2^)+(0.14*δ_N_*^2^) where Δδ is the weighted average chemical shift perturbation, Δδ_H_ is the change in ^1^H chemical shift, Δδ_N_ is the change in ^15^N chemical shift, and 0.14 is the weighting factor for ^15^N shifts.^41,42^

The apparent dissociation constants (*K*_D_) for binding of the Abz-LPATGGK(Dnp) substrate were determined from NMR chemical shift perturbation titration data by nonlinear curve fitting, assuming a 1:1 protein-ligand binding equilibrium. For selected amino acid resonances, the experimentally measured CSPs at each substrate concentration were fitted against CSP values calculated from the standard quadratic binding equation as previously described, according to the following equation: Δδ_cal_ = Δδ_max_ (*K*_D_ + [*P*]_t_ + [*L*]_t_ – {(*K*_D_ + [*P*]_t_ + [*L*]_t_)2 – (4 •[*P*]_t_ •[*L*]_t_)}1/ 2) /2[*P*]_t_.^41,43^ Here, Δδ_cal_ is the calculated CSP at a given ligand concentration, Δδ_max_ is the fitted maximum CSP at saturation, [*P*]_t_ is the total protein concentration, and [*L*]_t_ is the total ligand concentration. The values of *K*_D_ and Δδ_max_ were obtained by minimizing the difference between the experimentally measured and calculated CSP values across the titration series. The resulting residue-specific *K*_D_ values from the selected amino acid resonances were then averaged to obtain an overall estimate of the apparent *K*_D_.

### Structural analyses

AlphaFold3 was used for structural modeling.^44^ PyMOL (Schrödinger Software) was used for structural analyses and figure preparation.

## Results and Discussion

### Mutation of Y187 in saSrtA to interrogate P94-Y187 interaction

In our previous work, we characterized 18 variants of P94X saSrtA (excluding P94C due to the nearby catalytic Cys).^31^ We found that when tested against 4 peptide substrates varying only at the P2 position (LP**A**TG, LP**E**TG, LP**K**TG, LP**S**TG), most variants were more active than WT saSrtA.^31^ Our enzyme kinetics assays in combination with AlphaFold3 structural modeling suggested that substrate recognition (affecting *K*_m_ values) was sensitive to the relative surface charge of the peptide binding site. In the WT enzyme, the highly conserved R216 amino acid creates a positively charged surface that we reasoned would moderately bias the preference for a P2 Glu (LP**E**TG).^31^ Mutations at P94 affect the relative charged surface of this binding site, which indeed resulted in differential reactivity toward LP(A/E/K/S)TG substrates for different P94X variants.^31^ However, our results did not directly assess the role of Y187, which potentially forms contacts with residues at the P94 position (**Figure 1D**).

To investigate the P94-Y187 interaction from the β7-β8 loop, where Y187 is located, we characterized Y187A, Y187D, and Y187R saSrtA variants. Protein expression, purification, and peptide cleavage assays were conducted as previously reported, and as described in the Materials and Methods.^10–12,29,31,35,36^ Briefly, peptide substrates were labeled with an N-terminal Abz fluorophore and C-terminal Dnp quencher, as K(Dnp), as described in the Materials and Methods. For simplicity, we will refer to Abz-LP**A**TGGK(Dnp) as LP**A**TG moving forward. The only position varied was P2 (in **bold**). Abz was excited at λ = 320 nm, and fluorescence was monitored at λ = 420 nm following sortase-mediated cleavage between the P1 Thr and P1’ Gly positions.

We first tested LP**A**TG with our Y187X saSrtA variants (**Figure 2A**). The Y187R mutation was the only variant tested that increased relative activity. As normalized to WT saSrtA at t = 20 min, Y187R was ∼2-fold more active. This was compared to 0.5-fold for Y187A and 0.05-fold for Y187D, which was effectively inactive (**Figure 2B**). These trends were consistent with additional substrates (LP**X**TG, **X** = **E, K, S**), in that the Y187R mutation boosted activity over WT in all experiments. Furthermore, the Y187A mutation was consistently reduced as compared to WT, and the Y187D mutation showed little to no activity independent of substrate sequence (**Figure S1**). Although the structural basis for these effects remains unclear, it is apparent that substitutions at both positions, P94 and Y187, can substantially modulate saSrtA activity.

**Figure 2.**
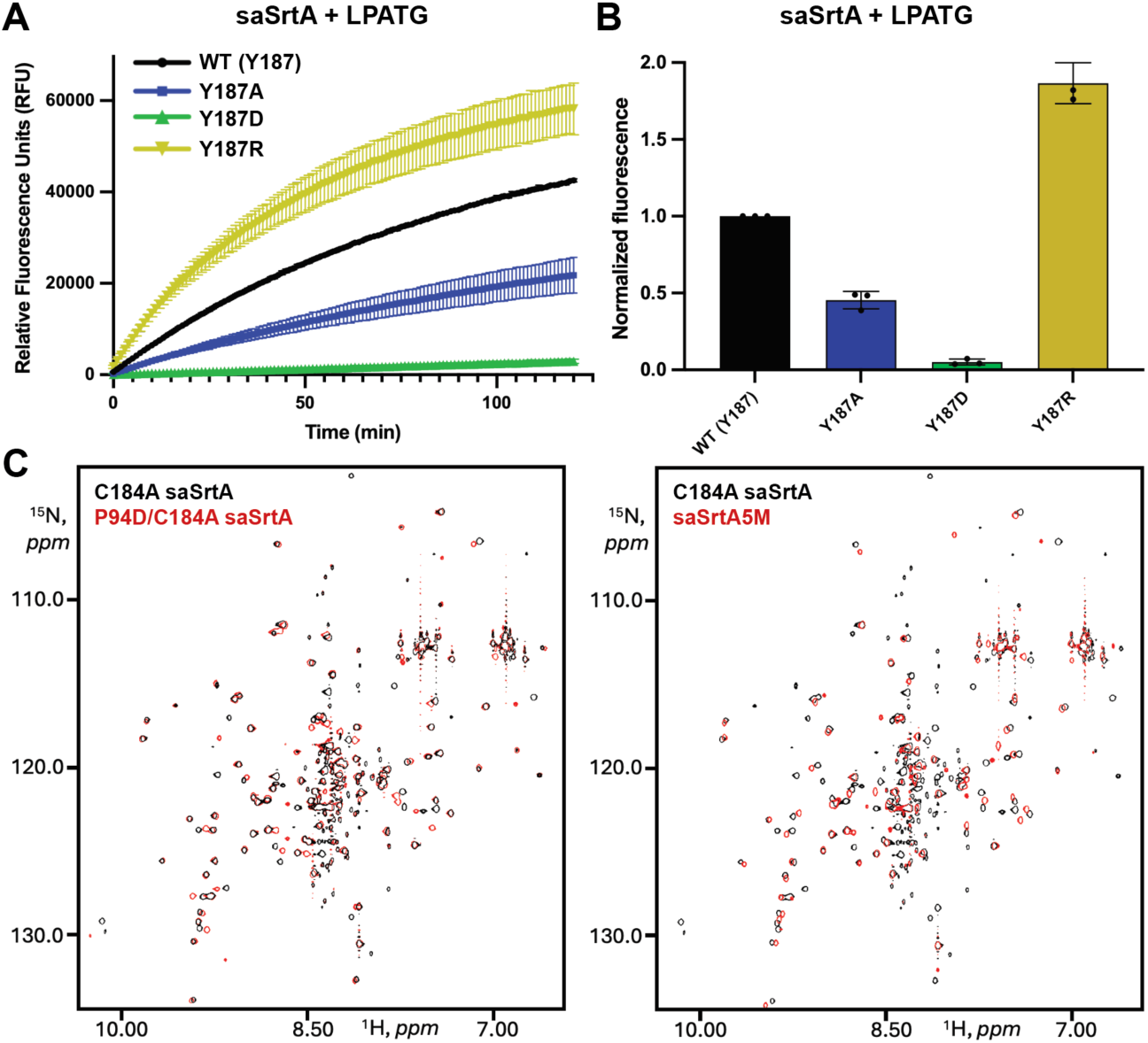
Peptide cleavage data for Y187X saSrtA variants and ^1^H-^15^N HSQC NMR spectra for inactive enzymes. (**A-B**) Peptide cleavage assay for Y187X saSrtA variants and the Abz-LPATGGK(Dnp) peptide substrate, as labeled. Data is the average from triplicate replicates, with standard deviation error bars shown. The full 2 h time course is in (**A**) and normalized data in (**B**), where the relative fluorescence units (RFU) for each variant at t = 20 min were normalized to WT saSrtA. (**C**) ^1^H-^15^N-HSQC NMR spectra for P94D/C184A saSrtA compared to C184A saSrtA (left) or saSrtA5M compared to C184A saSrtA (right). The peaks are colored as labeled.

### NMR and kinetic analyses of LPATG peptide binding

Early structural information of class A sortase enzymes was determined using NMR.^26,27^ More recently, NMR was used to determine the 3D structure of the transient acyl-enzyme intermediate.^13^ Therefore, we used NMR to further interrogate the effect of sortase mutation on recognition of a substrate peptide. To calculate binding affinities for the LP**A**TG peptide substrate, we expressed and purified ^15^N-labeled catalytically inactive C184A saSrtA, P94D/C184A saSrtA, and C184A saSrtA5M enzymes, as described in the Materials and Methods. We then collected ^1^H-^15^N heteronuclear single-quantum coherence (HSQC) NMR spectra in the presence of calcium and increasing concentrations of our LPATG peptide substrate. Spectra of 148-residue apo C184A and P94D/C184A saSrtA were largely similar to WT and each other with around 110 backbone HN signals being either unperturbed or perturbed minimally between C184A and P94D/C184A saSrtA samples (**Figure 2C, left**). Contrary to that, C184A and saSrtA5M varied more notably from each other (**Figure 2C, right**) with fewer than 70 unperturbed or weakly perturbed backbone signals. As a result, we were able to assign most resonances for C184A and P94D/C184A saSrtA by transferring assignments previously reported for the WT enzyme.^26,40^ Despite the spectral differences for C184A saSrtA5M, there were several unperturbed peaks that we were able to unambiguously identify, and these were used to monitor shifts during our substrate titration experiments.

To determine substrate binding affinities, here, as a *K*_D_ value due to the C184A inactivating mutation, we selected resonances that could be confidently assigned for each enzyme and which also showed chemical shift perturbations with increasing substrate concentration (**Figure 3**). For C184A saSrtA, these were F122, D124, R125, V161, V168, Y187, and G192. For P94D/C184A saSrtA, these were V168, N188, G192, and V193. For C184A saSrtA5M, these were A104, G119, K145, K173, E190, T191, and G192. We then determined the combined chemical shift perturbation (Δθ ^1^H-^15^N in ppm) for each individual residue, and averaged fits of these to calculate *K*_D_ values (**Figure 3**). We analyzed only those resonances that produced a clear plateau in their chemical shift trajectory in the *K*_D_ calculation. A small subset of resonances with strong perturbations (highlighted in **Figure 3A**) produced much more linear trajectories, rather than the hyperbolic behavior expected of saturation binding. It is possible that these sites represent a subset of residues distant from the binding site or a region of structure responding to a longer-range conformational change. Calculated *K*_D_ values for our variants with the LP**A**TG peptide substrate were 10 mM for C184A, 0.4 mM for P94D/C184A saSrtA, and 0.5 mM for C184A saSrtA5M. Here, we observed a 20-fold reduction between C184A saSrtA and C184A saSrtA5M, respectively, which fell within the range of previously reported values for *K*_m_ determined using a LP**E**TG substrate. Specifically, in our previously work we found a 4-fold reduction (*K*_m_ = 0.185 mM (WT) versus 0.047 mM (saSrtA5M)), while others have reported a 33-fold reduction (*K*_m_ = 7.6 mM (WT) versus 0.23 mM (saSrtA5M)).^16,31^ The similar *K*_D_ values for P94D/C184A saSrtA and C184A saSrtA5M were also consistent with our recently reported enzyme kinetics data for LP**E**TG, with *K*_m_ = 0.037 mM for P94D saSrtA and 0.047 mM for saSrtA5M.^31^

**Figure 3.**
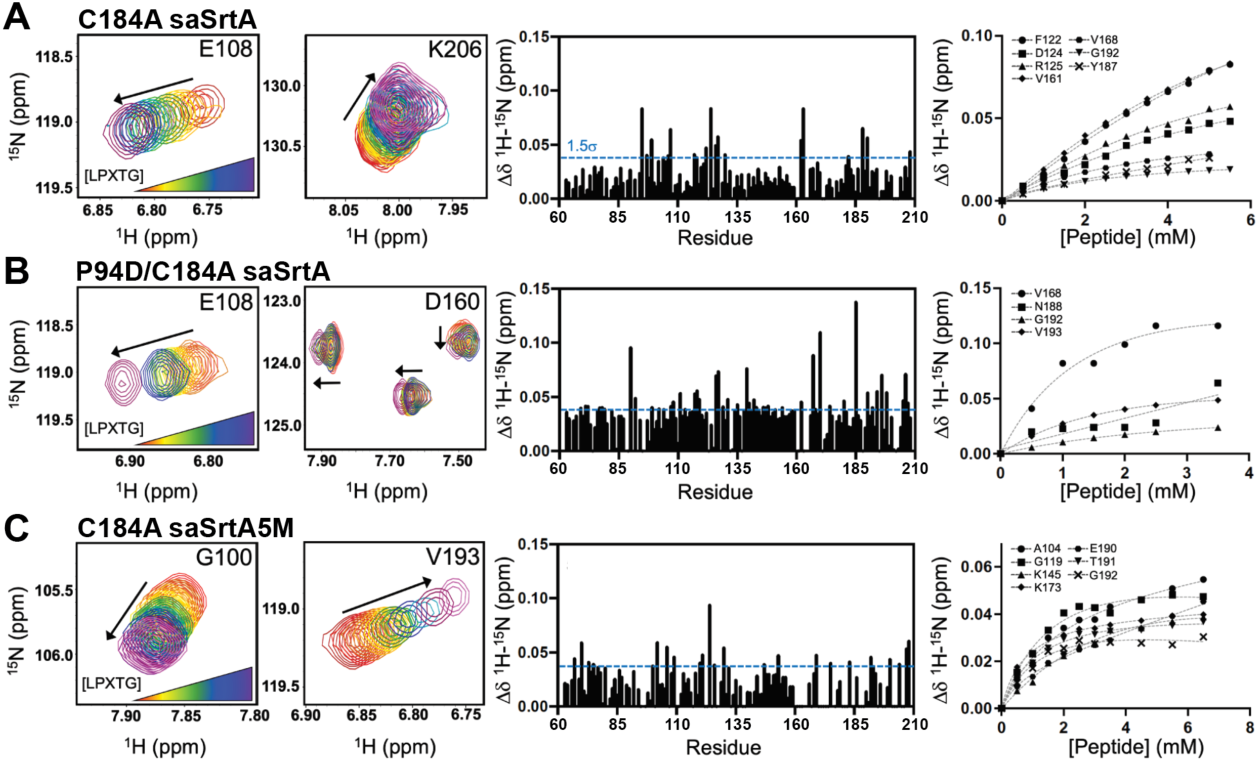
^1^H-^15^N HSQC experiments used to determine binding affinities of inactive saSrtA variants through titrations of LPATG peptide substrate. Example titration data (left), changes in CSPs graphed as a function of residue number (middle), and generated curves used to determine K_D_ values (right) are shown for C184A saSrtA (**A**), P94D/C184A saSrtA (**B**), and C184A saSrtA5M (**C**). Calculated binding affinities were K_D_ = 10 mM for C184A, 0.4 mM for P94D/C184A saSrtA, and 0.5 mM for C184A saSrtA5M.

To investigate these values further, we calculated *K*_m_ and *k*_cat_ for the active versions of WT and the P94D and saSrtA5M variants with the LP**A**TG peptide substrate (**Table 1**, **Figures 4, S2**).^31^ Determined values were consistent with our previous enzyme kinetics data for the LP**E**TG substrate, where *k*_cat_/*K*_m_ = 120 ± 1 M^-1^ s^-1^ for WT saSrtA, 550 ± 30 M^-1^ s^-1^ for P94D saSrtA, and 770 ± 120 M^-1^ s^-1^ for saSrtA5M.^31^ Interestingly, we observed a modest increase in *k*_cat_/*K*_m_ for the LP**A**TG substrate and P94D saSrtA (750 ± 60 M^-1^ s^-1^), such that it was equivalent with saSrtA5M (745 ± 85 M^-1^ s^-1^) (**Table 1**).^31^ This result is consistent with our determined *K*_D_ values from NMR (**Figure 3**) and our previous peptide cleavage assays that showed more similar enzyme activity for P94D saSrtA and saSrtA5M at t=20 min with LP**A**TG (RFU = 46944.7 ± 4129.8 and 59591.3 ± 1543.2, respectively, or P94D/saSrtA5M = 0.79) versus LP**E**TG (42246.7 ± 804.5 and 73917.7 ±1708.1, respectively, or P94D/saSrtA5M = 0.57). Taken together, these results support our previous conclusions that specificity for the P2 amino acid (LP**<u>X</u>**TG) is affected by the identity of the amino acid at position 94. Furthermore, the P94D saSrtA variant has similar activity to saSrtA5M with certain substrates, e.g., LP**A**TG (as well as LP**K**TG and LP**S**TG, based on our prior work).^31^ Finally, these data demonstrate the utility of *K*_D_ determination via solution NMR as a means to understand the activity of sortase variants, particularly those known to be driven largely by *K*_m_ differences.

**Figure 4.**
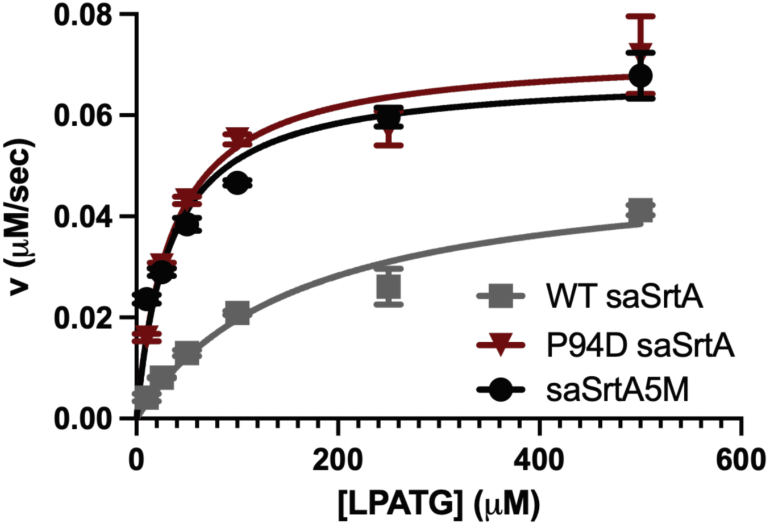
Enzyme kinetics assays of saSrtA variants with LPATG peptide substrate. The Michaelis-Menten curves for WT saSrtA (grey squares), P94D saSrtA (red triangles), and saSrtA5M (black circles) are shown. Data is the result of technical triplicate experiments, with standard deviation error bars. Calculated k_cat_ and K_m_ values are in **Table 1**.

**Table 1.** Kinetic characterization of saSrtA variants with the LPATG peptide substrate.

| | $k_{cat}$, S<sup>-1</sup> | $K_m$ LPATG, μM | $k_{cat}/K_m$ LPATG, M<sup>-1</sup> S<sup>-1</sup> |
| --- | --- | --- | --- |
| WT saSrtA | $0.018 \pm 0.001$ | $160 \pm 10$ | $115 \pm 10$ |
| P94D saSrtA | $0.026 \pm 0.001$ | $35 \pm 5$ | $750 \pm 60$ |
| P94R/D160N/D165A/K190E/K190T (saSrtA5M) | $0.024 \pm 0.001$ | $33 \pm 5$ | $745 \pm 85$ |

### Conformational differences in active apo saSrtA enzymes

Next, we used NMR data to quantify structural differences in the active apo states for the C184A and P94D/C184A saSrtA enzymes. Here, we assumed that any structural differences between these variants would be similar for the WT and P94D enzymes without the added inactivating C184A mutation. Comparison to data using the C184A saSrtA5M enzyme was excluded due to the lesser degree of unambiguous amino acid NMR resonance assignments. A full investigation of structural dynamics across timescales was also beyond the scope of this work, but we reasoned that our ^1^H-^15^N-HSQC spectra in the presence of Ca^2+^ and the absence of LP**A**TG could provide insight into whether amino acids distal or proximal to the P94D mutation showed marked NMR chemical shift variations.

While most resonances were minimally perturbed between C184A and P94D/C184A saSrtA, there are at least 25 peaks that differed substantially (**Figure 2C**) as determined by chemical shift perturbations (CSPs) > 0.04 ppm in the ^1^H and/or > 0.2 ppm in the ^15^N dimension(s) or line broadening in the P94D/C184A spectra. For this assessment, we excluded any resonances affected by spectral overlap or difficulty in transferring NMR assignments from BMRB entry 4879 from our analyses.^26,40^ We focused on peaks which corresponded to residues in the β7-β8 loop as we reasoned these were most likely to be affected by P94 mutation. Indeed, we observed several differences between the ^1^H-^15^N HSQC spectra of saSrtA variants in this region, including for D186, Y187, N188, V193, and W194. For example, while present in the C184A saSrtA spectra, the D186, Y187, and N188 signals in the P94D/C184A spectrum could not be identified reliably due to either line broadening or major CSPs (**Figure 5A**). The V193 and W194 resonances appeared to be present and identifiable in both C184A and P94D/C184A spectra, but clearly shifted with significant CSP between the two variants (**Figure 5B**).

**Figure 5.**
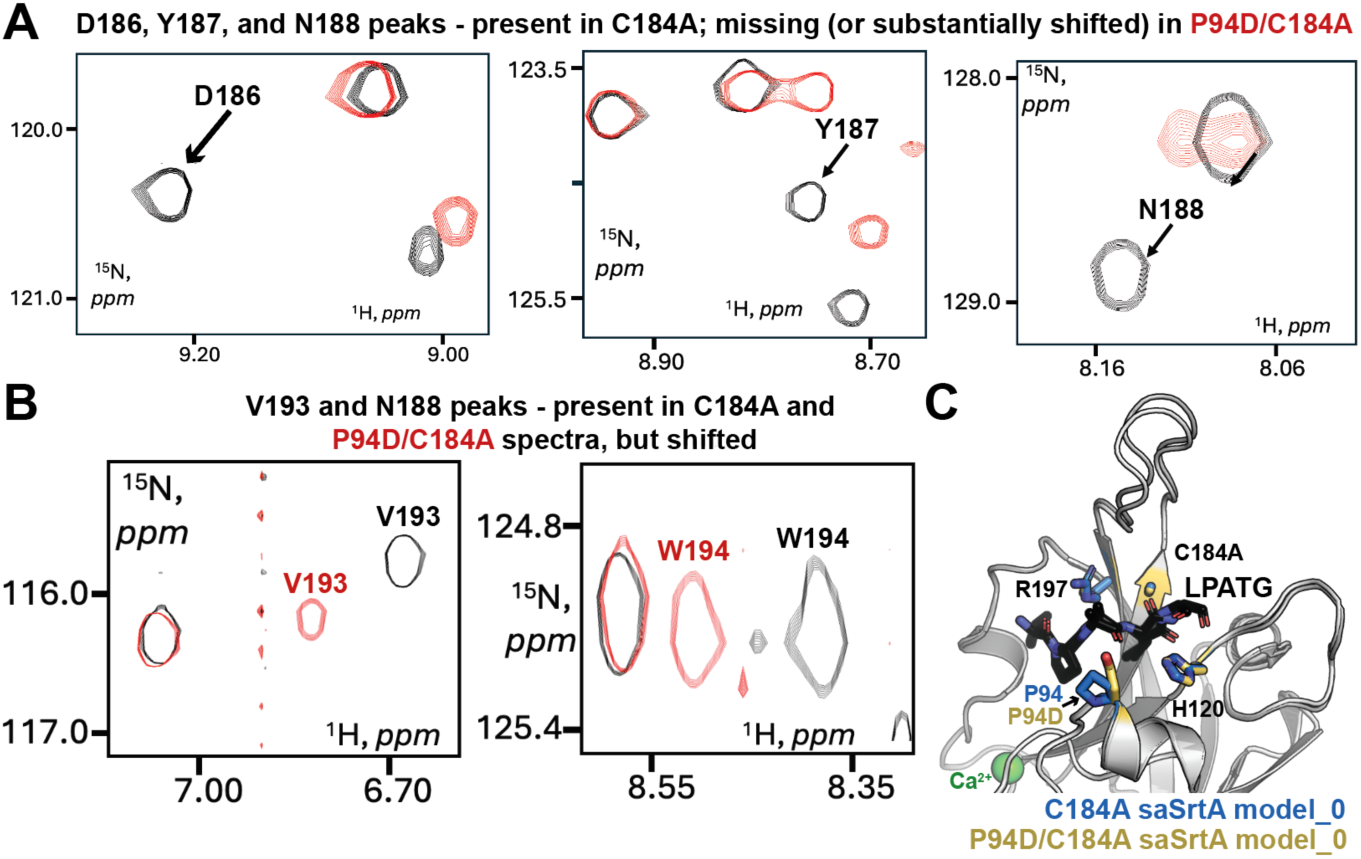
Differences in ^1^H-^15^N HSQC for C184A and P94D/C184A saSrtA in the presence of Ca^2+^ and absence of substrate, plus AlphaFold3 models of active, substrate-bound enzymes. (**A, B**) Differences in ^1^H-^15^N HSQC spectra for C184A and P94D/C184A saSrtA, + Ca^2+^, -substrate, are highlighted and labeled. These include peaks representing amino acids in the β7-β8 loop that are present in the C184A saSrtA spectrum but cannot be assigned in the P94D/C184A saSrtA spectrum because they are (**A**) broadened or very significantly shifted or (**B**) able to be assigned but shifted. (**C**) The AlphaFold3 output_0 models of C184A and P94D/C184A saSrtA, + Ca^2+^ + substrate are shown in cartoon representation, with the active site residues and P94 or P94D position shown as side chain sticks and colored by heteroatom as labeled (C=blue or yellow, O=red, N=blue). The Ca^2+^ is shown as a green sphere. The overall RMSD for these models is 0.134 Å (896 atoms).

We used AlphaFold3 to model the LPATG substrate-bound complexes of C184A and P94D/C184A saSrtA in the presence of Ca^2+^ (**Figures 5C, S3**). Because AlphaFold3 does not reliably model just a pentapeptide with saSrtA, we instead used <u>AQA</u>LPATGG, which includes the AQA sequence from an endogenous substrate, SpaA (UniProt ID: SPA_STAAN). Moreover, saSrtA is unlikely to bind an isolated LPXTG pentapeptide, as this motif would be present in an extended cell wall sorting signal (CWSS). Overall, good confidence Alphafold3 models were obtained which showed the AQALPETGG motif bound in the expected conformation (for C184A saSrtA-LPATGG + Ca^2+^, pTM =0.91 and ipTM = 0.73 and for P94D/C184A saSrtA-LPATGG + Ca^2+^, pTM = 0.89 and ipTM = 0.62) (**Figure S3A**). Consistent with the lower confidence of the latter, 2 of the 5 output models for P94D/C184A failed to position the substrate in its canonical binding site (**Figure S3B**), in which the P4 Leu occupies a hydrophobic binding pocket and the mutated A184 residue is proximal to the P1 Thr carbonyl carbon, the site of nucleophilic attack by the catalytic Cys.^2,3,12^

Comparison of the highest ranked (model_0) models revealed strong agreement in the overall structure (**Figure 5C**), with root-mean-squared-deviation (RMSD) = 0.134 Å over 896 atoms. Since the β7-β8 loop would be open in a substrate-bound state, and therefore the predicted P94D-Y187 interaction would be absent, this similarity was expected. However, we pursued this analysis because our ^1^H-^15^N HSQC data highlighted potential differences in the substrate-bound spectra as well. For example, the D186, Y187 and W194 resonances that broadened or underwent major CSPs in the spectra of apo P94D/C184A saSrtA remained absent or significantly shifted in the peptide titration (**Figures 6A, S4A, S4C**). Unlike D186 and Y197, the V193 signal for C184A saSrtA was unperturbed by the ligand titration and was observed in spectra of P94D/C184A saSrtA with the signal position and relative strength being sensitive to the ligand concentration (**Figure 6B**). In C184A saSrtA, the N188 signal weakened and eventually disappeared upon introduction of the ligand (**Figure S4B**). Overall, our NMR data reveal that the P94D mutation alters the dynamic properties of the β7-β8 loop, even in the substrate-bound state.

**Figure 6.**
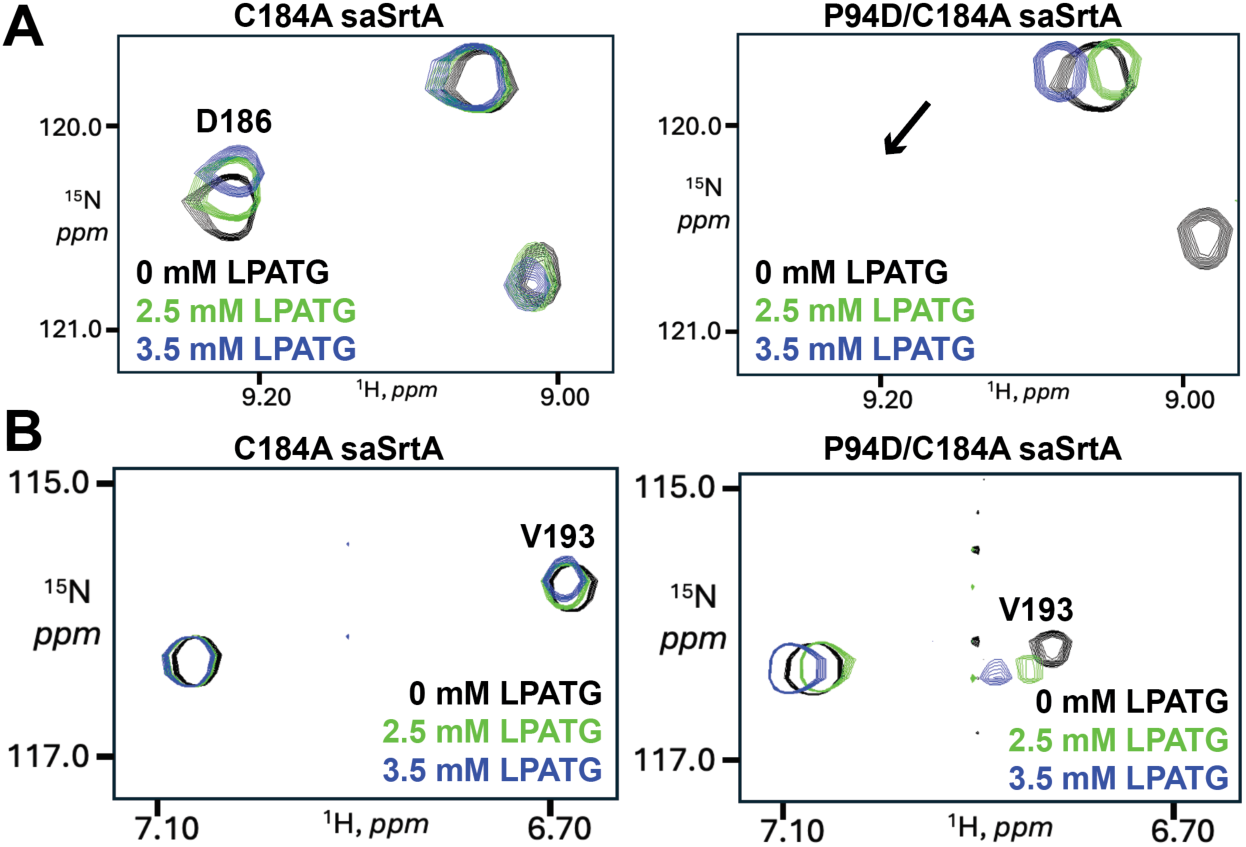
^1^H-^15^N HSQC data of select amino acid residues in presence of increasing substrate concentration for C184A and P94D/C184A saSrtA. (**A**) Comparison ^1^H-^15^N HSQC spectra for C184A (left) and P94D/C184A (right) saSrtA, highlighting the region surrounding the D186 amino acid peak for C184A saSrtA. Data in the presence of increasing substrate (LPATG peptide) concentration is colored as labeled. The arrow on the right indicates the area where the D186 peak is expected, but not present. (**B**) Comparison spectra (as in **A**) for the V193 amino acid, which is present and identifiable for both variants, but with differing spectral properties: upon introduction of P94D mutation, signals are shifting and changing relative strength (peak volumes).

## Conclusions

Experimental structures of class A sortases suggest distinct conformational dynamics in the β7-β8 loops of SrtA enzymes, as we previously reported.^12,26,27,31,45,46^ This includes a disordered-to-ordered transition for *Bacillus anthracis* SrtA upon substrate binding, in contrast to very little change in conformation in *Streptococcus pyogenes* SrtA.^31^ In saSrtA, experimental apo and ligand-bound NMR structures showed two distinct conformational ensembles, with one containing the β7-β8 loop in a closed, inactive position, that transitions upon substrate-binding to an open, active state.^31^ It remains unclear how conformational dynamics play a role in saSrtA substrate availability, e.g., the degree that the open, active β7-β8 loop conformation is sampled in the WT or amongst different saSrtA variants. A recent X-ray crystal structure of saSrtA5M in the apo +Ca^2+^, -substrate state (PDB ID 9WYA)^32^ revealed the open, active conformation for saSrtA5M in the presence of Ca^2+^, identical to our AlphaFold3 models with Ca^2+^ and substrate (**Figure 7**). Conformational dynamics in the β7-β8 loop clearly play a role in saSrtA catalytic efficiency, but questions remain about overall enzyme flexibility in the +Ca^2+^, apo state.

**Figure 7.**
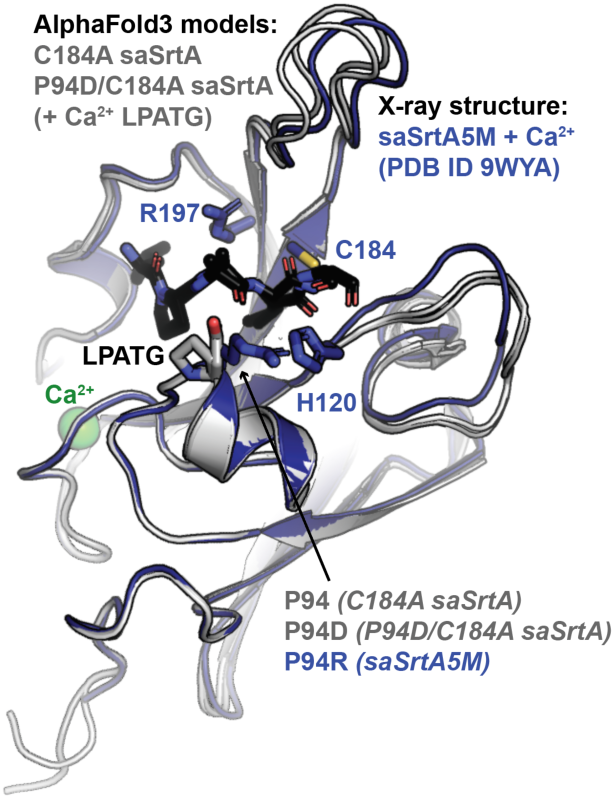
Alignment of experimental saSrtA5M crystal structure with AlphaFold3 models of active, substrate-bound C184A and P94D/C184A saSrtA. The AlphaFold3 models (C184A saSrtA and P94D/C184A saSrtA, with Ca^2+^ and substrate) and experimental saSrtA5M + Ca^2+^crystal structure (PDB ID 9WYA) are shown in cartoon representation. The structures are colored as labeled, with the side chains of the active site residues shown for saSrtA5M and colored by heteroatom (C=blue, N=blue, S=gold). The side chain atoms at the P94 position are also shown as labeled. The structures are very similar, with RMSD for saSrtA5M aligned to C184A = 0.446 Å (862 atoms) and 0.406 Å (866 atoms) for saSrtA5M aligned to P94D/C184A. The LPATG substrate in the AlphaFold3 models is shown as sticks and colored by heteroatom (C=black, O=red, N=blue). Calcium is shown as a green sphere and labeled.

Our overarching hypothesis is that disruption of the P94-Y187 interaction via mutation of P94, e.g., to Asp in P94D, facilitates the conformational transition to the open, active state of saSrtA. We argue that this allows for more favorable access of substrate to the active site and reduces the *K*_m_ of the interaction. This drives an increased catalytic efficiency of the enzyme, which is supported by the data presented here; for the LPATG substrate sequence, the *k*_cat_ and *K*_m_ were almost identical for P94D saSrtA and saSrtA5M, which contains P94R plus an additional 4 mutations (**Table 1**). Furthermore, ^1^H-^15^N HSQC NMR spectra revealed differences in several resonances corresponding to β7-β8 loop residues in the presence of the P94D mutation. We also showed that NMR can be used to estimate relative binding affinities for catalytically inactive saSrtA variants. Future studies characterizing the specific dynamics for the β7-β8 loops between these variants, as well as completing resonance assignments for saSrtA5M, will shed more light on the structural basis for sortase specificity. Taken together, this study builds upon our previous work and provides additional insight into substrate recognition by this workhorse enzyme that can be applied to next-generation SML technologies.

## Supporting information

Supplemental Information for Walkenhauer et al.

## Author Contributions

**Erich G. Walkenhauer**: Investigation (lead), Formal analysis (supporting), Writing – Review & Editing (supporting), **Noah Cox-Tigre**: Investigation (lead), Formal Analysis (supporting), Writing – Review & Editing (supporting), **Manish Chaubey**: Investigation (supporting), Formal Analysis (supporting), Visualization (supporting), Writing – Review & Editing (supporting), **Raphael Marcenac**: Formal Analysis (supporting), Writing – Review & Editing (supporting), **Adam Wachsman**: Investigation (supporting), Writing – Review & Editing (supporting), **Hanna M. Kodama**: Conceptualization (supporting), Writing – Review & Editing (supporting), **Katy M. Lindblom**: Investigation (supporting), **Claire E. Bloom**: Investigation (supporting),**John M. Antos**: Formal Analysis (supporting), Supervision (supporting), Writing – Review & Editing (supporting), **George P. Lisi**: Visualization (supporting), Formal Analysis (supporting), Supervision (supporting), Funding Acquisition (supporting), Resources (supporting), Writing – Review & Editing (supporting), **Serge L. Smirnov**: Visualization (lead), Formal Analysis (supporting), Visualization (supporting), Supervision (supporting), Resources (supporting), Writing – Review & Editing (supporting), **Jeanine F. Amacher**: Conceptualization (lead), Visualization (supporting), Resources (lead), Funding Acquisition (lead), Supervision (lead), Writing – Original Draft Preparation (lead)

## Acknowledgements

The authors want to sincerely thank all members of the Amacher lab for research support and useful discussion, specifically to C. Ceravolo for assistance with structural analyses. This work was supported by NSF CHE-2044958, a Cottrell Scholar Award from the Research Corporation for Science Advancement, and a Henry Dreyfus Teacher-Scholar Award from the Camille and Henry Dreyfus Foundation to J.F. Amacher. Additional funding was provided to E.G. Walkenhauer and A. Wachsman by Western Washington University Research and Sponsored Programs. A. Wachsman was also funded by NSF REU CHE-2243968. N. Cox-Tigre was supported by a Beckman Scholars Award from the Arnold and Mabel Beckman Foundation. G.P. Lisi acknowledges support from NSF grant MCB-2143760. Preliminary NMR data acquisition was performed on a Bruker Avance III HD NMR spectrometer (500 MHz ^1^H frequency) at Western Washington University, which was acquired via NSF MRI CHE-1532269 grant, awarded to S.L. Smirnov.

