## Supplemental Information for Walkenhauer et al. for "Mutation-induced heterogeneity of the β7-β8 loop of the *Staphylococcus aureus* class A sortase leading to enhanced catalytic efficiency characterized by NMR and enzyme kinetics"

**Table of Contents**

**Figure S1. Peptide cleavage data for Y187X saSrtA variants with different substrates.** 2

**Figure S2. Replicate enzyme kinetics data for saSrtA variants and the LPATG peptide substrate.** 3

**Figure S3. AlphaFold3 models of saSrtA with $Ca^{2+}$ and LPATG substrate.** 4

**Figure S4. $^{1}H$-$^{15}N$ HSQC data of select amino acid residues in presence of increasing substrate concentration for C184A and P94D/C184A saSrtA.** 5

**Sequences used in this study.** 6

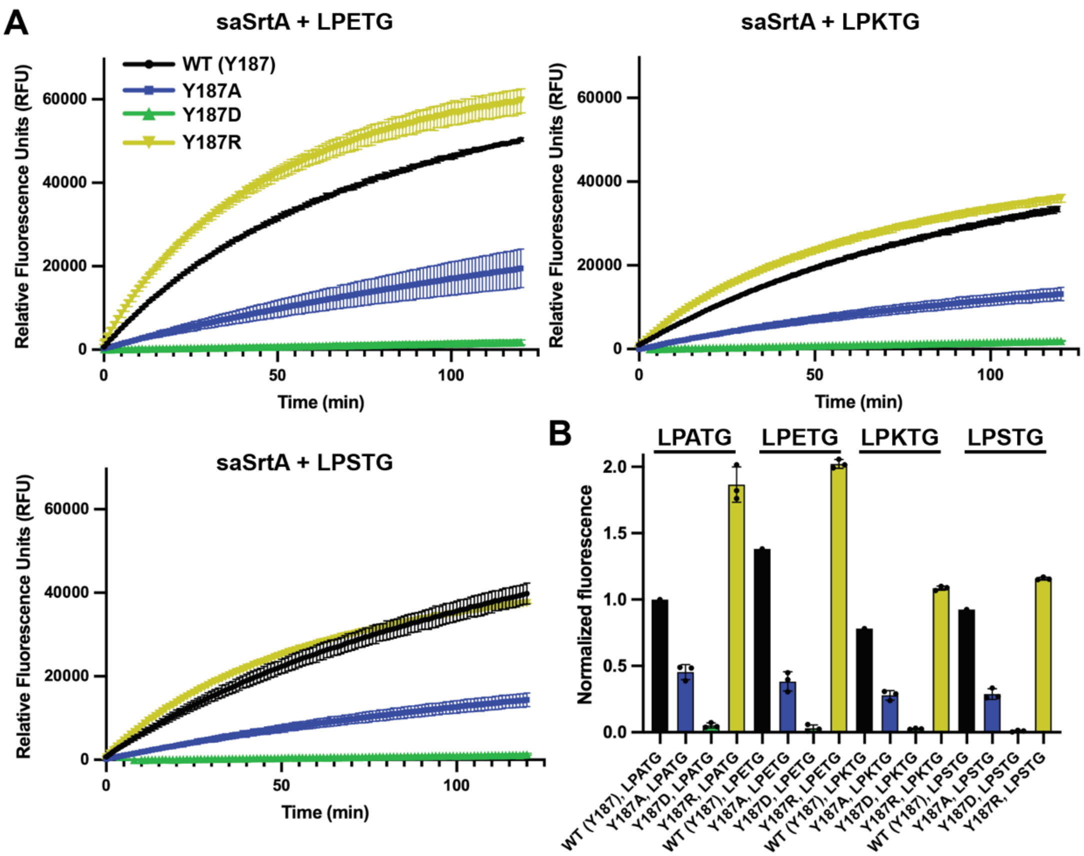


**Figure S1. Peptide cleavage data for Y187X saSrtA variants with different substrates.** (**A**) Cleavage assay time course data over t = 2 h for Y187X variants and the Abz-LP**E**TGGK(Dnp), Abz-LP**K**TGGK(Dnp), and Abz-LP**S**TGGK(Dnp) peptide substrates. Averaged data from technical triplicate replicates are shown, with standard deviation error bars. (**B**) The data at t = 20 min was normalized to the relative fluorescence units (RFU) for WT saSrtA with the Abz-LP**A**TGGK(Dnp) substrate.

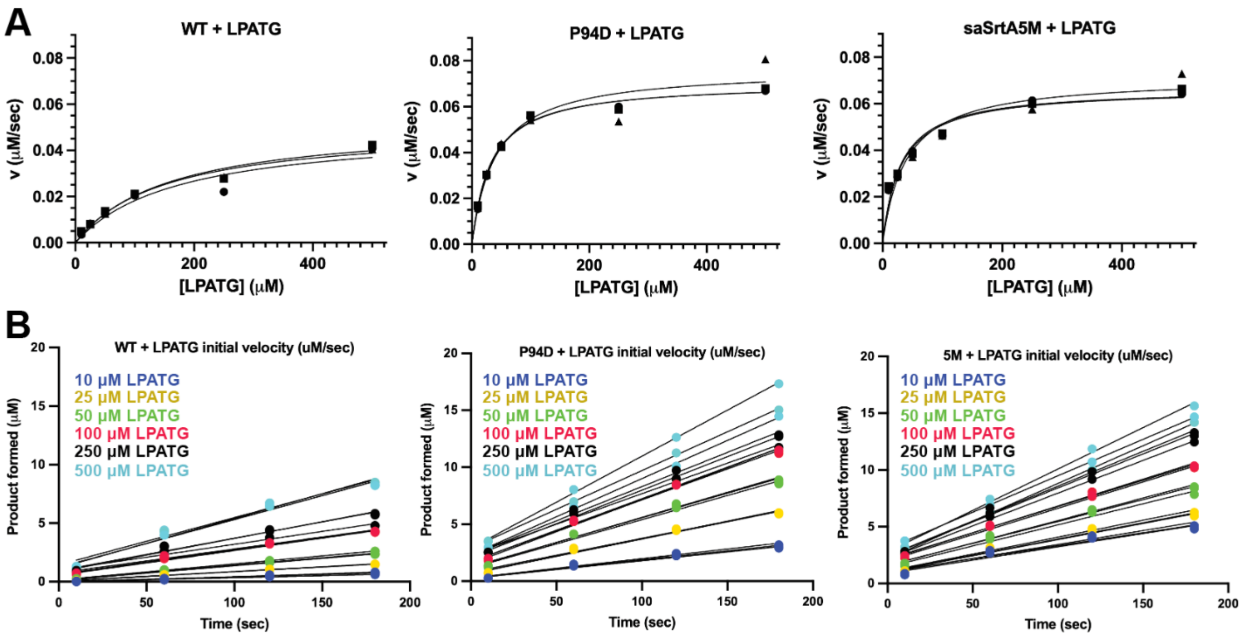


**Figure S2. Replicate enzyme kinetics data for saSrtA variants and the LPATG peptide substrate.** (**A**) Replicate Michaelis-Menten curves are shown for WT saSrtA, P94D saSrtA, and saSrtA5M with the Abz-LP**A**TGGK(Dnp) peptide substrate. Initial velocities were calculated following integration of the product peak using an HPLC assay over time. Assays were conducted in triplicate. (**B**) Replicate product integration data (in μM) are shown for the peptide titrations with each saSrtA variant. Each replicate is colored according to the peptide concentration used, as labeled in the key. A linear regression curve was calculated for each replicate, with the slope equal to initial velocity, v (μM/sec).

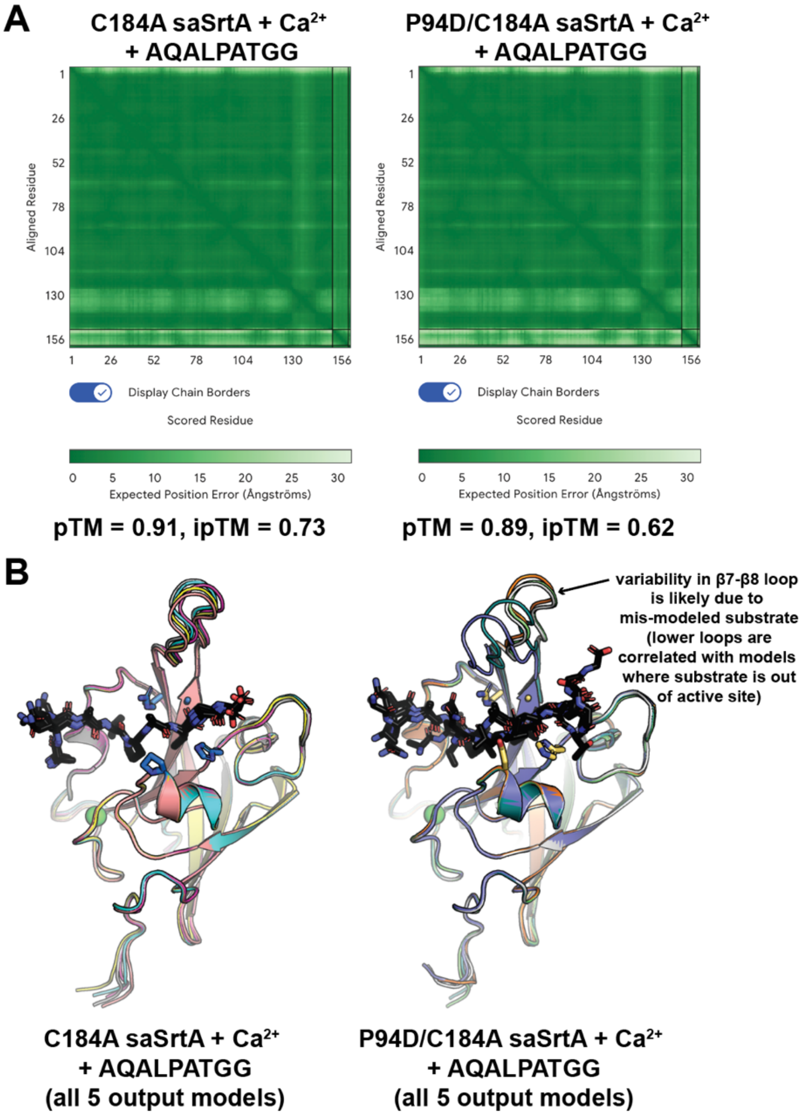


**Figure S3. AlphaFold3 models of saSrtA with $Ca^{2+}$ and LPATG substrate.** (**A**) AlphaFold3 output data for the models calculated. (**B**) All 5 output models are shown for C184A saSrtA + $Ca^{2+}$ + AQALPATGG (left) and P94D/C184A saSrtA + $Ca^{2+}$ + AQALPATGG (right) in cartoon representation. The peptides are in sticks and colored by heteroatom (C=black, O=red, N=blue). The side chain atoms for active site residues and the P94 (or P94D) position are shown for model_0 and colored by heteroatom (C=marine/yellow, O=red, N=blue). The calcium ion is shown as a green sphere. The peptide was not properly positioned for multiple output models in the P94D/C184A model (right), which likely affected the position of the β7-β8 loop, i.e., it was shifted closer to the down, inactive conformation. Structural analysis of the peptide position with respect to the active site C184A, as well as positioning of the P4 Leu, suggested model_0 was properly modeled; thus, model_0 was used for comparison with C184A saSrtA.

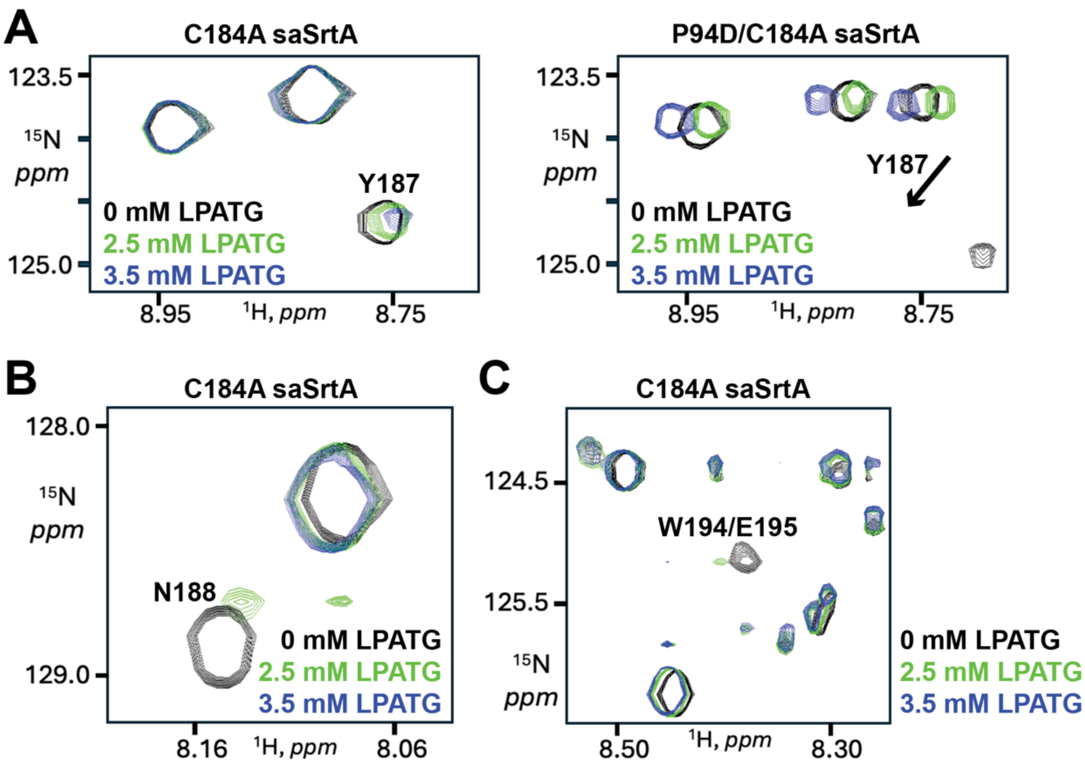


**Figure S4. $^1$H-$^{15}$N HSQC data of select amino acid residues in presence of increasing substrate concentration for C184A and P94D/C184A saSrtA.** (**A**) Comparison $^1$H-$^{15}$N HSQC spectra for C184A (left) and P94D/C184A (right) saSrtA, highlighting the region surrounding the Y187 amino acid peak for C184A saSrtA. Data in the presence of increasing substrate (LPATG peptide) concentration is colored as labeled. The arrow on the right indicates the area where the Y187 peak is expected, but not present. (**B**) Titration data for C184A saSrtA shows that while there is a peak for N188 in the absence of peptide, there are no observable peaks for N188 with higher concentrations of LPATG. This contrasts with the amino acid peak nearby, which shows signal for the higher peptide concentrations. (**C**) Spectra for W194, which are present only for C184A saSrtA (as in **B**). In SrtA WT, the backbone $^1$H and $^{15}$N resonance assignments for W194 are nearly identical to those for an adjacent residue E195 (BMRB entry 4879) thus we labeled the signal as originating from both residues. Both N188 and W194/E195 resonances are detectible in the apo form of C184A SrtA and quickly diminish in strength upon introduction of the LPATG ligand.

**Sequences used in this study.** Underlined amino acids indicate the 6xHis tag used for protein purification and TEV protease cleavage site (ENLYFQ/S, where "/" is the cleavage site). Bold indicates differences from the WT saSrtA sequence. All sequences were inserted into the pET28a(+) expression plasmid. The WT saSrtA sequence matches UniProt ID SRTA_STAA8.

>saSrtA
MESSHHHHHHENLYFQSQAKPQIPKDKSKVAGYIEIPDADIKEPVYPGPATPEQLNRGVSFAEENESLDDQNISIAG
HTFIDRPNYQFTNLKAAKKGSMVYFKVGNETRKYKMTSIRDVKPTDVGVLDEQKGKDKQLTLITCDDYNEKTGVWEK
RKIFVATEVK

>C184A_saSrtA
MESSHHHHHHENLYFQSQAKPQIPKDKSKVAGYIEIPDADIKEPVYPGPATPEQLNRGVSFAEENESLDDQNISIAG
HTFIDRPNYQFTNLKAAKKGSMVYFKVGNETRKYKMTSIRDVKPTDVGVLDEQKGKDKQLTLIT**A**DDYNEKTGVWEK
RKIFVATEVK

>P94D_saSrtA
MESSHHHHHHENLYFQSQAKPQIPKDKSKVAGYIEIPDADIKEPVYPGPAT**D**EQLNRGVSFAEENESLDDQNISIAG
HTFIDRPNYQFTNLKAAKKGSMVYFKVGNETRKYKMTSIRDVKPTDVGVLDEQKGKDKQLTLITCDDYNEKTGVWEK
RKIFVATEVK

>P94D_C184A_saSrtA
MESSHHHHHHENLYFQSQAKPQIPKDKSKVAGYIEIPDADIKEPVYPGPAT**D**EQLNRGVSFAEENESLDDQNISIAG
HTFIDRPNYQFTNLKAAKKGSMVYFKVGNETRKYKMTSIRDVKPTDVGVLDEQKGKDKQLTLIT**A**DDYNEKTGVWEK
RKIFVATEVK

>Y187A_saSrtA
MESSHHHHHHENLYFQSQAKPQIPKDKSKVAGYIEIPDADIKEPVYPGPATPEQLNRGVSFAEENESLDDQNISIAG
HTFIDRPNYQFTNLKAAKKGSMVYFKVGNETRKYKMTSIRDVKPTDVGVLDEQKGKDKQLTLITCDD**A**NEKTGVWEK
RKIFVATEVK

>Y187D_saSrtA
MESSHHHHHHENLYFQSQAKPQIPKDKSKVAGYIEIPDADIKEPVYPGPATPEQLNRGVSFAEENESLDDQNISIAG
HTFIDRPNYQFTNLKAAKKGSMVYFKVGNETRKYKMTSIRDVKPTDVGVLDEQKGKDKQLTLITCDD**D**NEKTGVWEK
RKIFVATEVK

>Y187R_saSrtA
MESSHHHHHHENLYFQSQAKPQIPKDKSKVAGYIEIPDADIKEPVYPGPATPEQLNRGVSFAEENESLDDQNISIAG
HTFIDRPNYQFTNLKAAKKGSMVYFKVGNETRKYKMTSIRDVKPTDVGVLDEQKGKDKQLTLITCDD**R**NEKTGVWEK
RKIFVATEVK

>saSrtA5M
MESSHHHHHHENLYFQSQAKPQIPKDKSKVAGYIEIPDADIKEPVYPGPAT**R**EQLNRGVSFAEENESLDDQNISIAG
HTFIDRPNYQFTNLKAAKKGSMVYFKVGNETRKYKMTSIR**N**VKPT**A**VGVLDEQKGKDKQLTLITCDDYNE**E**TGVWE**T**
RKIFVATEVK

>C184A_saSrtA5M
MESSHHHHHHENLYFQSQAKPQIPKDKSKVAGYIEIPDADIKEPVYPGPAT**R**EQLNRGVSFAEENESLDDQNISIAG
HTFIDRPNYQFTNLKAAKKGSMVYFKVGNETRKYKMTSIR**N**VKPT**A**VGVLDEQKGKDKQLTLIT**A**DDYNE**E**TGVWE**T**
RKIFVATEVK
